## Supplementary information for "Selective lipid recruitment by an archaeal DPANN symbiont from its host"

### Supplementary Materials

### Supplementary Figures 1-8 and Discussion

### Supplementary Discussion

#### Growth Measurements

Optical density-based growth curves of pure *Hrr. lacusprofundi* cultures and co-cultures containing *Ca.* Nha. antarcticus indicated that co-cultures displayed slightly reduced growth rates relative to pure cultures. This effect is strongest in the 6 and 12 h timepoints suggesting that presence of the nanohaloarchaeon initially inhibits growth of much of the host cells in the culture before growth rate increases to similar levels as those in the pure cultures in the 24 and 48 h timepoints. The increased rate of growth coincides with the statistically significant increase in MK abundance within co-culture lipid biomass which is consistent with an upregulation of respiration and therefore energy conservation. Together these results suggest that initial introduction of *Ca.* Nha. antarcticus to *Hrr. lacusprofundi* causes a decreased growth rate until the host adjusts to the increased metabolic load of the symbiont at which point growth rates return to similar levels as those in pure *Hrr. lacusprofundi* cultures. This is consistent with previous studies of DPANN-host interactions that have reported an upregulation of metabolic processes and downregulation of proliferation and division processes within *Ignicoccus hospitalis* when interacting with *Nanoarchaeum equitans*[1].

qPCR data showed a stable 16s rRNA copy number for *Hrr. lacusprofundi* across all time points and in both culture conditions despite $OD_{600}$ values changing consistent with exponential growth. It has previously been established that haloarchaea are highly polyploid and that genome copy number is actively changed over the course of culture growth with highest copy number at mid-exponential and lowest at stationary phase[2]. The qPCR results observed for *Hrr. lacusprofundi* are consistent with a similar regulation of genome copy number whereby despite cells continuing to actively divide (as evidenced by increasing $OD_{600}$) replication is effectively halted and the existing genomes are distributed amongst daughter cells with progressively fewer copies. In contrast, *Ca.* Nha. antarcticus displayed an increase in 16s rRNA copy number between 24 and 48 h indicating large population growth consistent with our recently proposed predatory lifestyle for the nanohaloarchaeon[3].

#### Visualisation of Interactions

FISH microscopy was conducted at the 12, 24, and 48 h timepoints in order to visualise interactions between *Ca.* Nha. antarcticus and *Hrr. lacusprofundi* and to identify differences in host morphology and cell size between co-cultures and pure cultures. Quantitative data indicates that statistically significant differences in host morphology and cell size exist between all timepoints and conditions with the exception of size between the two 24 h timepoints (Supplementary Figure 5, Supplementary Table 1). Generally, despite statistical significance the trend in cell size is similar between both pure and co-cultures with a progressive decrease in average cell size that may reflect the culture beginning to enter nutrient limiting conditions and deceleration cellular proliferation processes. However, cell shape displayed a significant difference in trend with pure cultures suggesting increased circularity over the course of the incubation whilst co-cultures became more rod-shaped between 12 and 24 h before showing increased circularity at the 48 h timepoint. *Hrr. lacusprofundi* is known to be pleomorphic and belongs to a sister genus to that of *Haloferax volcanii* which has been shown to regulate cell shape as a function of growth stage[4]. The trend observed in pure cultures suggests *Hrr. lacusprofundi* may regulate morphology in a similar manner and the difference in cell shape in co-cultures may come as a consequence of the presence of *Ca.* Nha. antarcticus and its impact on the growth stage of co-cultures compared to pure cultures.

Several different stages of interactions could be discerned during co-culture growth including examples of *Ca.* Nha. antarcticus attached to the surface of *Hrr. lacusprofundi* (Supplementary Fig. S2) as well as instances where the two cells were indistinguishable which may be indicative of latter stages of predation (Supplementary Fig. S3). Consistent with previous reports *Ca.* Nha. antarcticus typically co-fluoresced for *Hrr. lacusprofundi* specific probes[5]. Interestingly, large cells that co-fluoresced for both organisms 16S rRNA probe frequently appeared to have DNA and RNA localised to different regions of the cell (Supplementary Fig. S4). Fluorescence intensity profiles reveal that this segregation of the two types of nucleic acids was not complete (as has been reported in Asgard archaea[6]) but show clear enrichment of signal from DNA and RNA to different locations within the cell with reduced signal in others. The reason for this distinct localisation is unclear: it may be induced by Ca. Nha. antarcticus as part of its predation of Hrr. lacusprofundi or it may be a response of

*Hrr. lacusprofundi* to the nanohaloarchaeon. Future work will be necessary to elucidate the processes behind the distinct localisation of DNA and RNA in *Hrr. lacusprofundi* cells interacting with *Ca.* Nha. antarcticus.

### References


1. Giannone RJ*, et al.* Life on the edge: functional genomic response of Ignicoccus hospitalis to the presence of Nanoarchaeum equitans. *Isme Journal* **9**, 101-114 (2015).

2. Breuert S, Allers T, Spohn G, Soppa J. Regulated polyploidy in halophilic archaea. *PLoS One* **1**, e92 (2006).

3. Hamm JN*, et al.* The parasitic lifestyle of an archaeal symbiont. *bioRxiv*, 2023.2002.2024.529834 (2023).

4. de Silva RT, Abdul-Halim MF, Pittrich DA, Brown HJ, Pohlschroder M, Duggin IG. Improved growth and morphological plasticity of Haloferax volcanii. *Microbiology (Reading)* **167**, (2021).

5. Hamm JN*, et al.* Unexpected host dependency of Antarctic Nanohaloarchaeota. *Proc Natl Acad Sci U S A* **116**, 14661-14670 (2019).

6. Avci B*, et al.* Spatial separation of ribosomes and DNA in Asgard archaeal cells. *ISME J* **16**, 606-610 (2022).

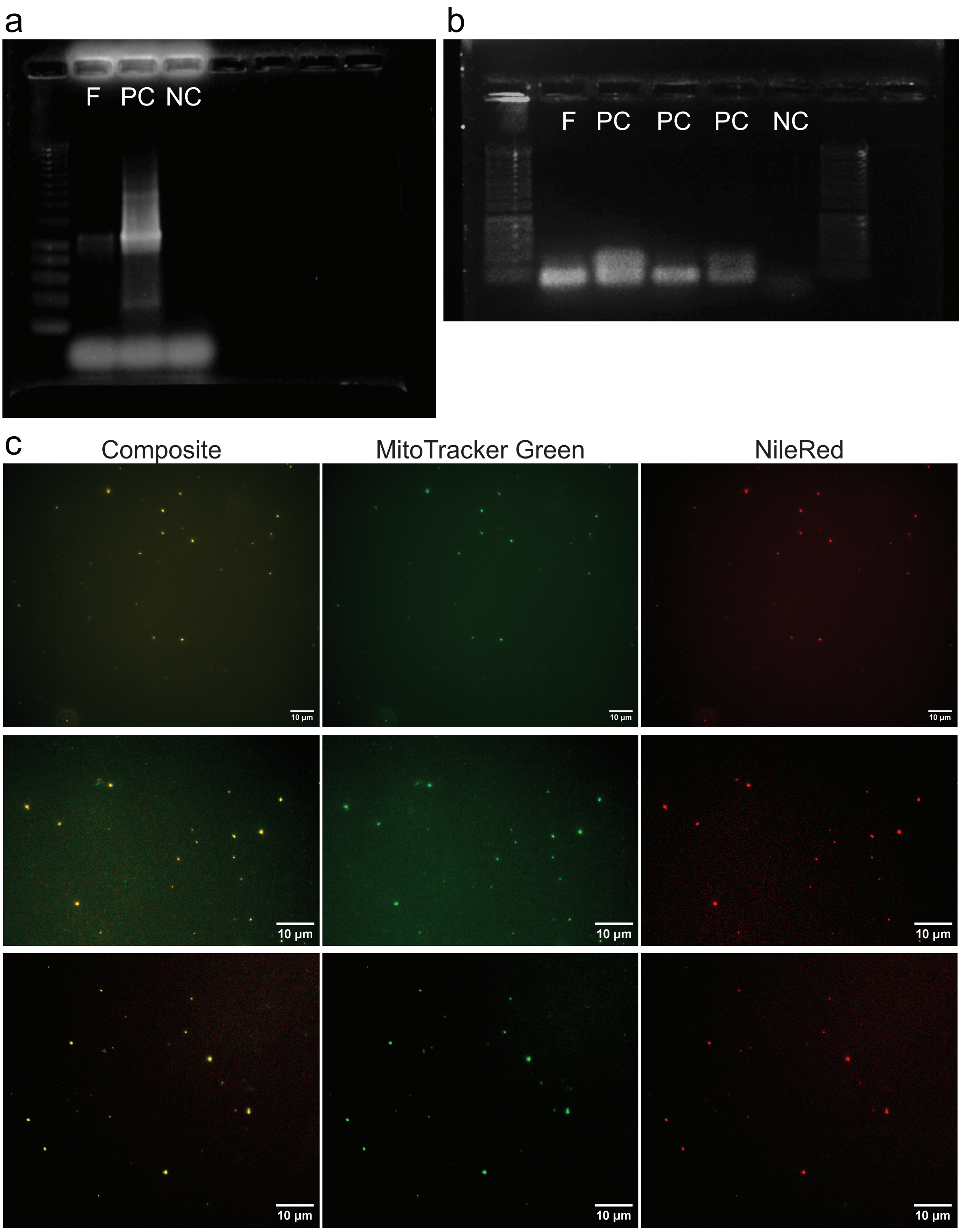


**Supplementary Fig. 1** a & b) Gel electrophoresis analysis of PCR products from filter purified *Ca.* Nha. antarcticus cells targeting either the a) *Hrr. lacusprofundi* 16S rRNA gene or b) *Ca.* Nha. antarcticus 16S rRNA gene. F: Filtered Cells, PC: Positive Control, NC: Negative Control. c) Fluorescence microscopy visualisation of filtered cells using MitoTracker Green (non-cytotoxic live cell stain) and Nile Red (lipid stain).

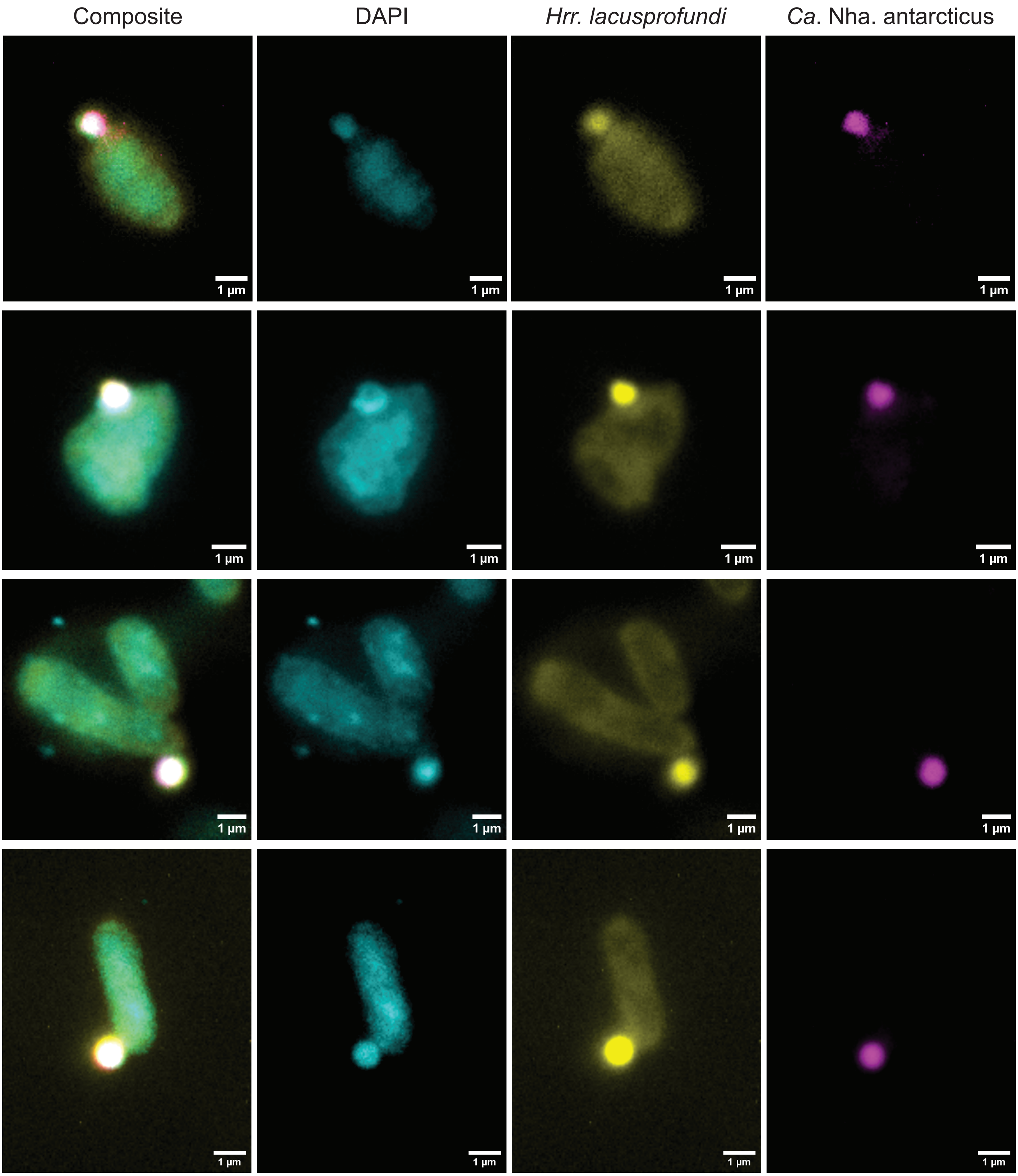


**Supplementary Fig. 2** FISH microscopy showing interactions between *Ca*. Nha. antarcticus and *Hrr*. *lacusprofundi* where the nanohaloarchaeon and host cell are still clearly distinguishable from each other. Images are consistent with early stages of the predation cycle previously proposed for *Ca*. Nha. antarcticus. Channels represent DNA (DAPI, Blue), *Hrr. lacusprofundi* 16S rRNA (*Hrr. lacusprofundi*, Yellow), and *Ca*. Nha. antarcticus 16S rRNA (*Ca*. Nha. antarcticus, Magenta).

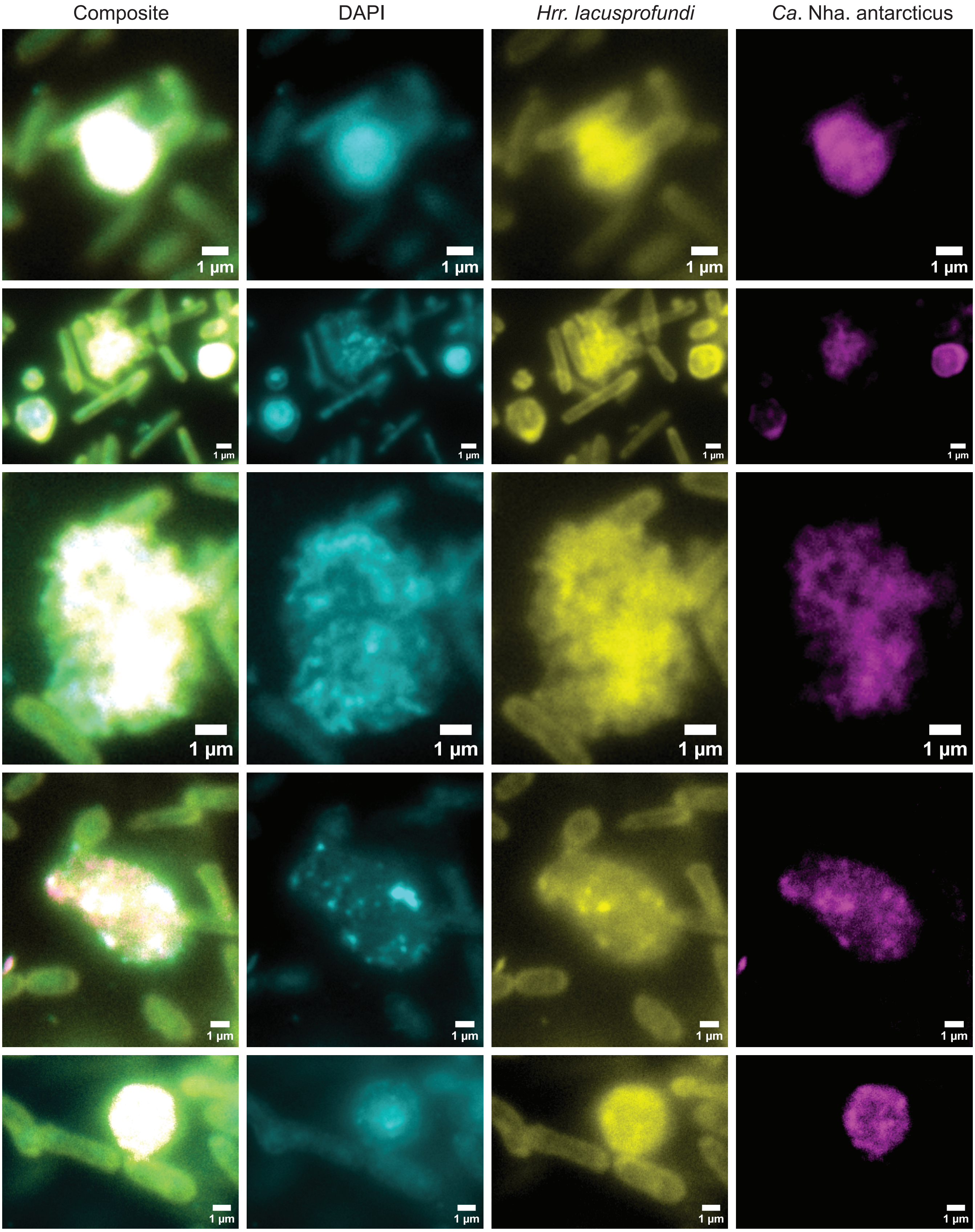


**Supplementary Fig. 3** FISH microscopy showing examples of interactions between *Ca*. Nha. antarcticus and *Hrr. lacusprofundi* where identites of the two cells cannot be distinguished. Images are consistent with late stages of the predation cycle previously proposed for *Ca*. Nha. antarcticus. Channels represent DNA (DAPI, Blue), *Hrr. lacusprofundi* 16S rRNA (*Hrr. lacusprofundi*, Yellow), and *Ca*. Nha. antarcticus 16S rRNA (*Ca*. Nha. antarcticus, Magenta).

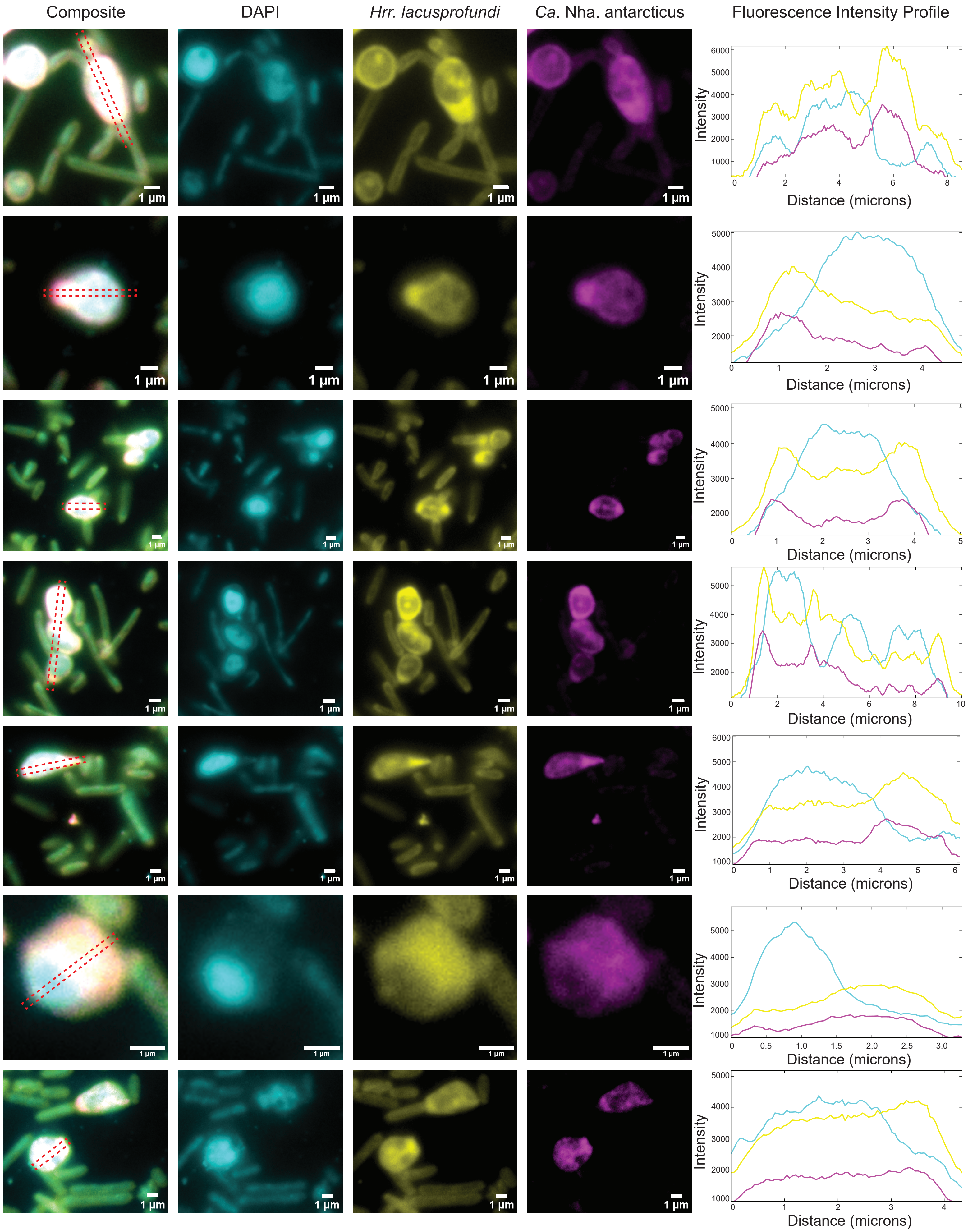


**Supplementary Fig. 4** FISH microscopy showing examples of interactions between *Ca*. Nha. antarcticus and *Hrr. lacusprofundi* where DNA and RNA appears to be localised to different regions of the cell. Channels represent DNA (DAPI, Blue), *Hrr. lacusprofundi* 16S rRNA (*Hrr. lacusprofundi*, Yellow), and *Ca*. Nha. antarcticus 16S rRNA (*Ca*. Nha. antarcticus, Magenta). The fluorescence intensity profile of all three channels along a cross-section of the cell in the composite (red dotted box) is shown.

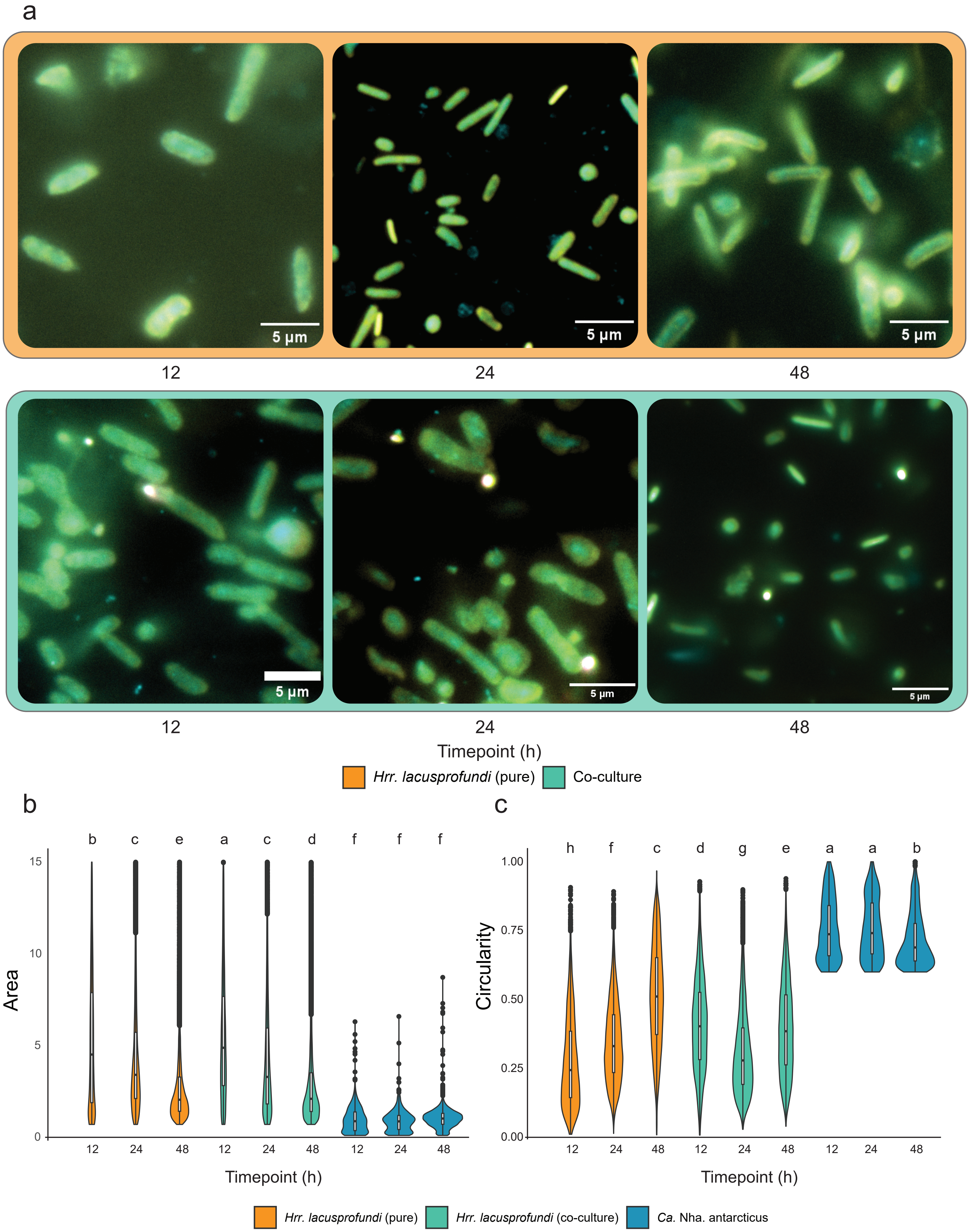


**Supplementary Fig. 5** Cell size and shape variation across cutivation conditions. a) Representative composite 16S rRNA FISH microscopy images of cultures at each timepoint sampled during incubation. Channels represent DNA (DAPI, Blue), *Hrr. lacusprofundi* 16S rRNA (*Hrr. lacusprofundi*, Yellow), and *Ca*. Nha. antarcticus 16S rRNA (*Ca*. Nha. antarcticus, Magenta). b and c) Violin plot showing variation in b) cell size and c) circularity of *Ca*. Nha. antarcticus and *Hrr. lacusprofundi* calculated from 16S rRNA images. Letters above plot indicate statistically significant variations as determined by Tukey's Honest Significance Difference Test (TukeyHSD) and visualised using Compact Letter Display (CLD) ($P < 0.05$).

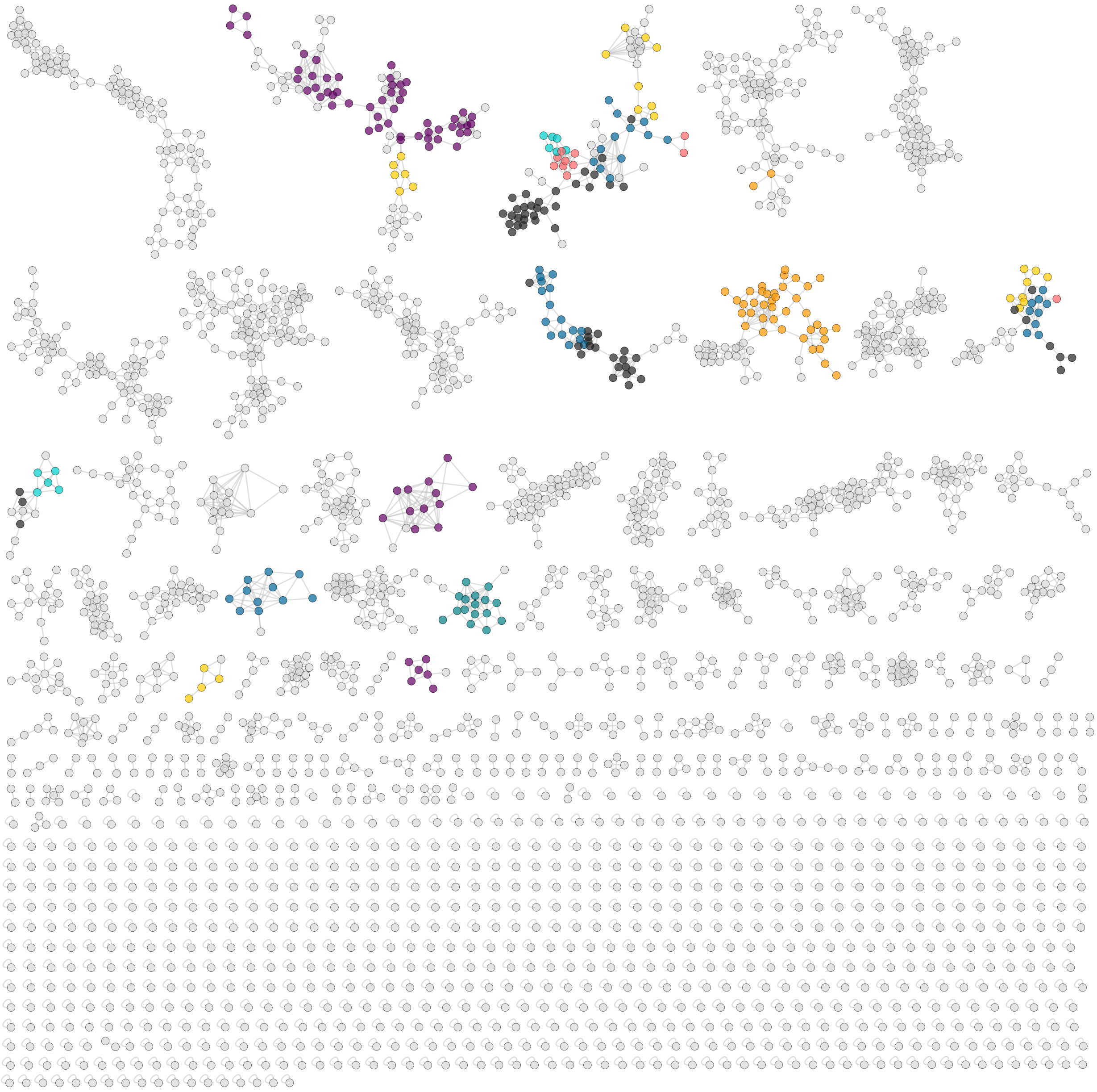


**Supplementary Fig. 6** The full view of molecular network of the *Hrr. lacusprofundi-Ca.* Nha. antarcticus system. Nodes represent $MS^2$ spectra of ion components (lipids), which are connected based on spectral similarity (cosine > 0.5). The spatial orientation of the nodes in the $MS^2$ network is randomly generated by Cytoscape[57, 58] and does not relate to relationships between the subnetworks. Lipid classes (clusters) with colors are either tentatively identified in this study or have been annotated from the GNPS library (Figs. 1d and 2, details in the Method Section).

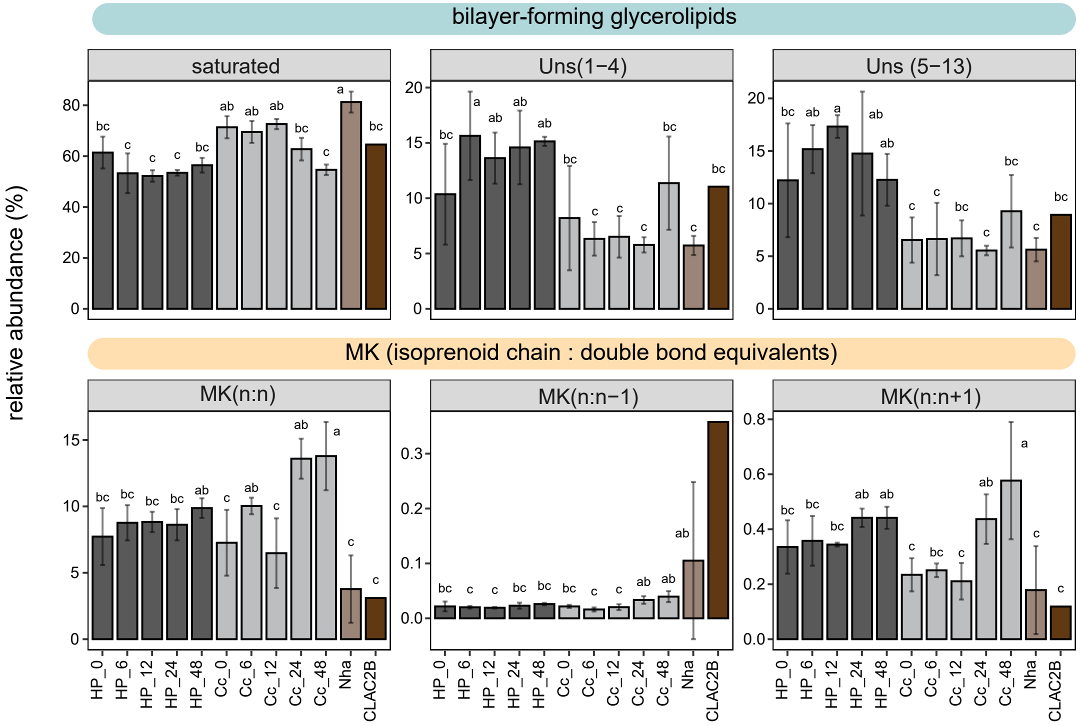


**Supplementary Fig. 7** The distribution of different unsaturation degree of the summed bilayer-forming glycerolipids and menaquinones among the samples. Statistical differences in lipid species among the samples were assessed using the Tukey's Honest Significance Difference test (TukeyHSD), and results were visualized with the Compact Letter Display (CLD) ($P < 0.05$).

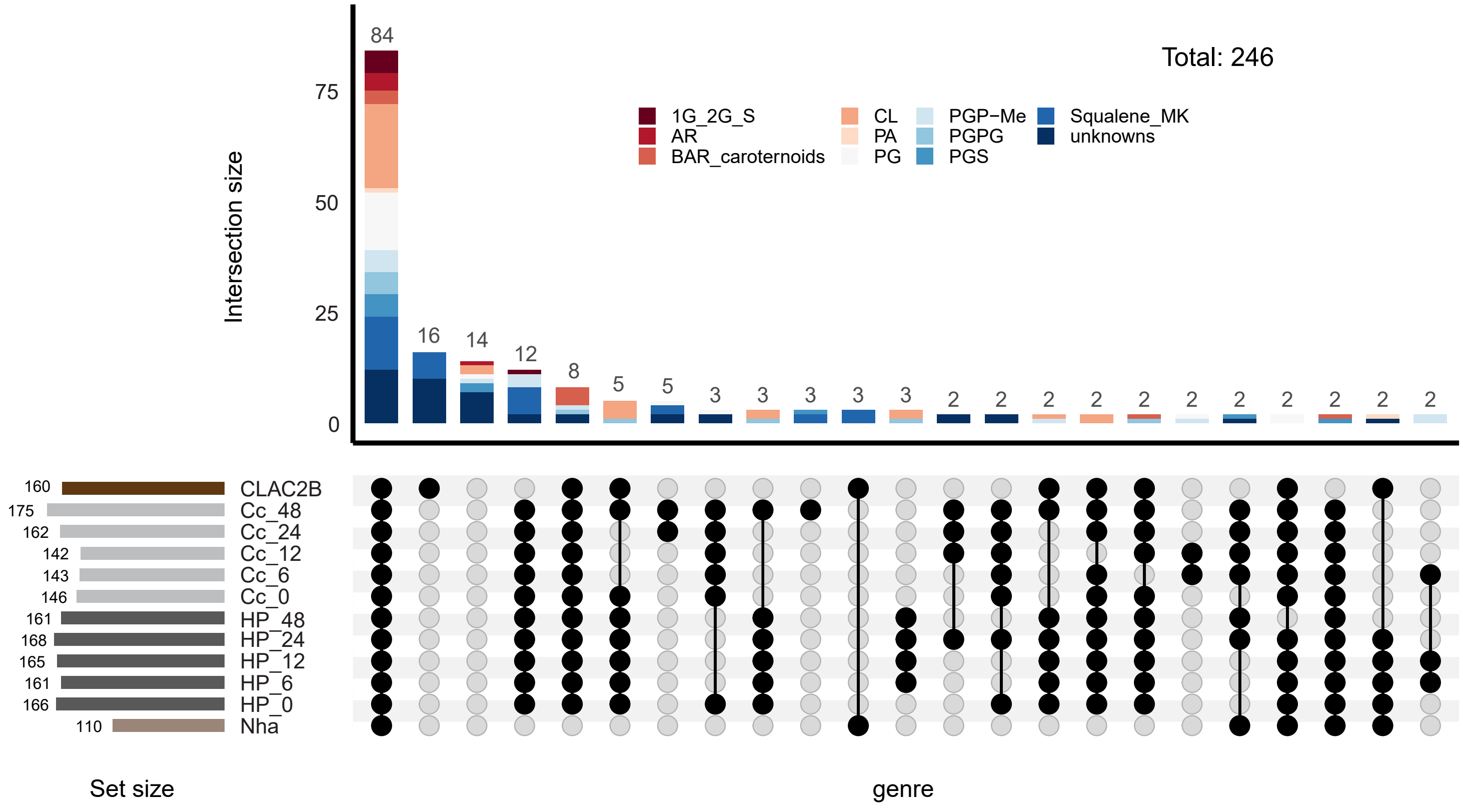


**Supplementary Fig. 8** The intersection of lipid species across samples including the enrichment is illustrated through an UpSet plot. A threshold of 0.01% relative abundance of total lipids was applied to determine the presence of a lipid in a specific sample; lipids with less than 0.01% of total lipid abundance were considered absent in that sample. The dark connected dots denote lipid species shared among these samples.
